# Frataxin deficiency induces lipid accumulation and affects thermogenesis in brown adipose tissue

**DOI:** 10.1101/664649

**Authors:** Riccardo Turchi, Flavia Tortolici, Giulio Guidobaldi, Federico Iacovelli, Mattia Falconi, Stefano Rufini, Raffaella Faraonio, Viviana Casagrande, Lorenzo De Angelis, Massimo Federici, Simone Carotti, Maria Francesconi, Maria Zingariello, Sergio Morini, Roberta Bernardini, Mattei Maurizio, Daniele Lettieri-Barbato, Katia Aquilano

## Abstract

Decreased expression of the mitochondrial protein frataxin (FXN) causes Friedreich’s ataxia (FRDA). FRDA is a neurodegenerative disease also characterized by systemic metabolic alterations that increase the risk of developing type 2 diabetes thus aggravating FRDA prognosis. Brown adipose tissue (BAT) is a mitochondria-enriched and anti-diabetic tissue that, in addition to its thermoregulatory role, turns excess energy into heat to maintain energy balance. Here we report that the FXN knock-in/knock-out (KIKO) mouse shows reduced energy expenditure and VO_2_, hyperlipidemia, decreased insulin sensitivity and enhanced circulating levels of leptin, recapitulating diabetes-like signatures. FXN deficiency leads to alteration of mitochondrial structure and oxygen consumption, decreased lipolysis and lipid accumulation in BAT. Transcriptomic data highlighted a blunted thermogenesis response, as several biological processes related to thermogenesis (e.g. response to temperature stimuli, mitochondrial gene transcription, triglyceride metabolism, adipogenesis) resulted affected in BAT of KIKO mice upon cold exposure. Decreased adaptation to cool temperature in association with limited PKA-mediated lipolysis and downregulation of the expression of the genes controlling mitochondrial metabolism and lipid catabolism were observed in KIKO mice. T37i brown adipocytes and primary adipocytes with FXN deficiency showed reduced thermogenesis and adipogenesis markers respectively recapitulating the molecular signatures detected in KIKO mice.

Collectively our data point to BAT dysfunction in FRDA and suggest BAT as a promising target to overcome metabolic complications in FRDA.

## INTRODUCTION

Friedreich’s Ataxia (FRDA) is an inherited autosomal recessive neurodegenerative disorder. It is caused by mutation in the gene encoding mitochondrial protein frataxin (FXN) (Abrahao et al., 2015). The primary function of FXN is to direct the mitochondrial synthesis of iron-sulfur clusters (Fe/S), which are essential parts of several mitochondrial enzymes, including mitochondrial respiratory chain complexes and aconitase (Maio and Rouault, 2015).

Deficiency in mitochondrial respiration, mitochondrial iron accumulation and oxidative stress are claimed as the main pathogenic factors in FRDA. Mitochondrial dysfunction mostly affects heart (Koeppen et al., 2015) and cerebellum at the level of dentate nucleus (Koeppen et al., 2011;Koeppen and Mazurkiewicz, 2013).

FRDA is characterized by a variable phenotype. Besides neurological symptoms, cardiomyopathy (Koeppen et al., 2015) and systemic metabolic alterations occur (Cnop et al., 2013), which can predispose to diabetes development and cause premature death. In particular, patients with FRDA experience a greater risk of abnormal glucose homeostasis, in the form of both insulin resistance and glucose intolerance (Isaacs et al., 2016). Increased blood cholesterol and triglycerides levels have been observed in FRDA patients (Raman et al., 2011;Tamarit et al., 2016). To date, the major cause of diabetes mellitus occurrence in FRDA patients seems to be related to impairment of mitochondria that in pancreatic β-cell are fundamental in generating signals that trigger and amplify insulin secretion (Cnop et al., 2012).

Brown adipose tissue (BAT) is a high oxidative tissue extremely rich in mitochondria that highly expresses the thermogenic protein uncoupling protein 1 (UCP1). BAT has emerged as a key regulator of glucose, lipid and insulin metabolism (Chondronikola et al., 2014;Chondronikola et al., 2016;Hankir and Klingenspor, 2018). Actually, BAT activity requires the uptake of substrates from the circulation, mostly free fatty acids (FAs), but also glucose, successfully leading to hypolipidemic and hypoglycemic effects which can significantly improve insulin sensitivity and exert a protective role in the pathogenesis of type 2 diabetes (Sidossis and Kajimura, 2015). FAs liberated from intracellular TGs through the action of lipases are also critical for BAT thermogenesis (Blondin et al., 2017). Interestingly, abnormal accumulation of intracellular lipids has been observed in patients’ cells as well as in animal models (Puccio et al., 2001;Coppola et al., 2009;Tamarit et al., 2016;Stram et al., 2017), pointing to a possible inefficient lipolysis.

Importantly, it has been discovered that decreased activity of BAT is associated with insulin resistance and diabetes (Flachs et al., 2013;Sacks and Symonds, 2013). Despite this potential clinical importance, the regulation of BAT in FDRA patients is not well-investigated.

In this work we show that FXN deficiency significantly affects lipolytic and thermogenic pathways as well as thermogenic adipocyte differentiation suggesting that BAT impairment could be at center stage of type 2 diabetes development in FRDA patients.

## RESULTS

### Alteration of basal metabolic parameters in KIKO mice

Herein, we used KIKO mouse that represents a suitable *in vivo* model for studying neurodegeneration as well as metabolic complications in FRDA (Coppola et al., 2009;McMackin et al., 2017). We firstly monitored body weight and a trend to gain weight was observed in 6-months KIKO mice that becomes significant at 8 months of age (**Fig. 1A**). In contrast, food and water intake were never found changed (**data not shown**). We have performed bio-clinical analyses and, among the tested parameters, a significant increase of triglycerides and cholesterol levels was detected in KIKO mice both at 6- and 8-months of age (**Fig. 1B**). Other metabolic parameters such as fasting glycaemia (**Fig. 1B**) as well as markers of general organ functions (total plasma proteins), including kidney (creatinine, urea) and liver (albumin) resulted unaltered (**data not shown**), suggesting that hyperlipidemia represents an early event in FRDA. Even though fasting glycaemia appeared unaffected, oral glucose tolerance test (OGTT) carried out in 8 months old KIKO mice revealed that glycaemia remained higher than WT mice at 120 min from glucose administration (**Fig 1C**).

**Fig. 1.**
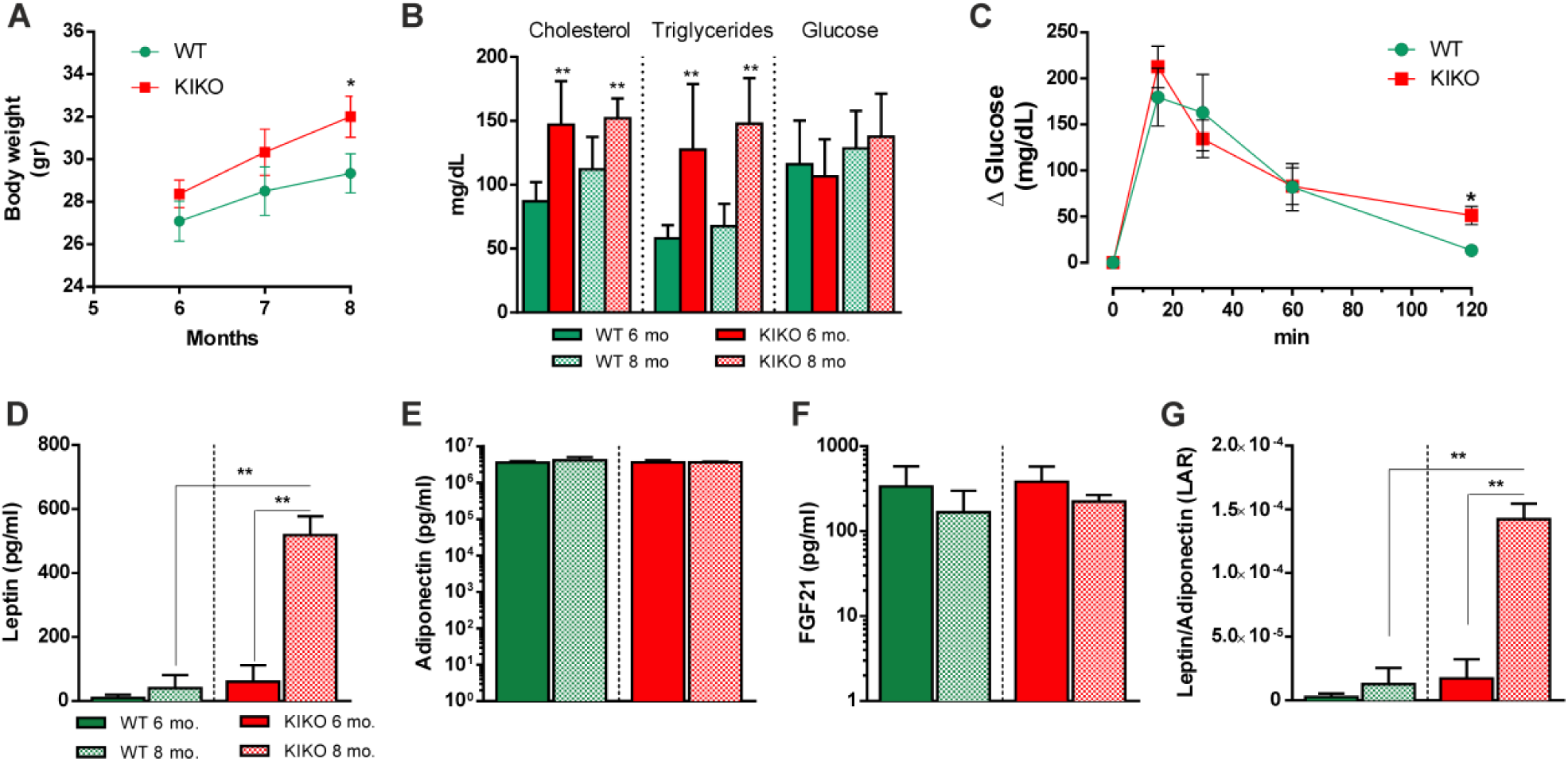
Bio-clinical and adipokine analyses revealed dyslipidemia and insulin resistance in KIKO mice. **(A)** Body weight recorded at different ages (n=12 each group; *p<0.05 vs age-matched WT mice). **(B)** Bio-clinical analyses of fasting glycaemia and lipidemia (triglycerides and cholesterol) carried out at different ages (n=12 each group; **p<0.001 vs age-matched WT mice). **(C)** Oral glucose tolerance test (OGTT) carried out in 8-months old mice (n=6 each group; *p<0.05 vs WT mice). (**D-G**) Serum levels of leptin (D), adiponectin (E), FGF21 (F) and leptin to adiponectin ratio (G) determined in mice at different ages (n=6 each group; **p<0.01).

The circulating levels of adipose tissue-secreted cytokines, i.e. adipokines, nicely reflect body metabolic state and are considered valid markers to monitor insulin resistance (Li et al., 2013;Adams-Huet et al., 2014). Through Luminex^®^ Multiplex Assay, we found that KIKO mice underwent a prominent raise of plasma leptin levels at 8 months of age (**Fig. 1D**), while other adipokines such as adiponectin and FGF21 remained unchanged (**Fig. 1E** and **1F**). The increase in leptin, and more specifically the leptin to adiponectin ratio (LAR), is considered a useful tool to evaluate insulin resistance (Finucane et al., 2009). Accordingly, 8 months-old KIKO mice showed increased LAR compared to WT mice (**Fig. 1G**). We have then characterized metabolic profile of KIKO mice by performing indirect calorimetry. Before measurements, all mice were acclimatized for 48 h into individual metabolic chambers at 25 °C, with free access to food (standard diet) and water. The respiratory exchange ratio (RER), which is expressed as VCO_2_/VO_2_, was similar in WT and KIKO mice and around 1.0 indicating carbohydrates as the predominant fuel source (**data not shown**). At 6 months, the recorded oxygen consumption (VO_2_) was lower in KIKO than WT mice, with KIKO mice showing a further decrease of VO_2_ at 8 months (**Fig. 2A**). Moreover, KIKO mice showed a decrease of resting energy expenditure (REE) at 6 months of age (**Fig. 2B**). This parameter was altered at higher extent at 8 months of age pointing to a progressive dysfunction in basal BAT thermogenic capacity (**Fig. 2B**).

**Fig. 2.**
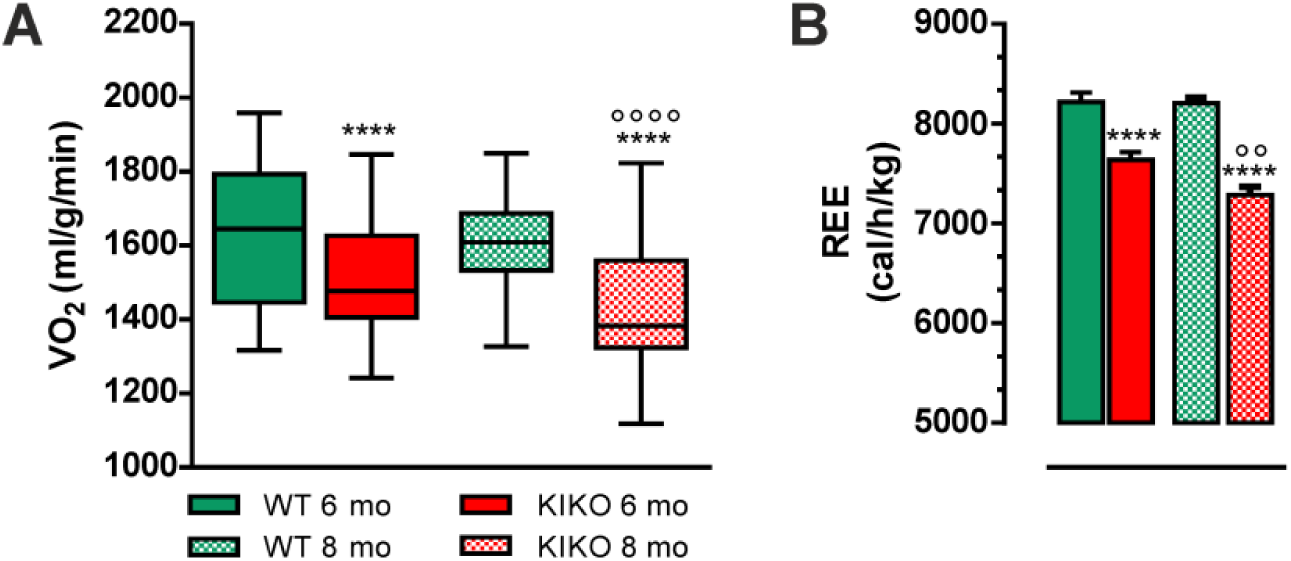
Indirect calorimetry indicated decreased oxygen consumption and energy expenditure in KIKO mice. (**A, B**) Oxygen consumption (VO_2_) (A) and resting energy expenditure (REE) (B) measured by indirect calorimetry at different ages (n=12; each group; ****p<0.0001 vs age-matched WT mice, °°°°p<0.0001,°°p<0.01 vs 6-months KIKO mice).

### KIKO mice show altered mitochondrial function and lipid accumulation in BAT

BAT activity strongly depends on mitochondrial lipid oxidation to produce heat (Hankir and Klingenspor, 2018). As dysfunctional FXN affects mitochondrial oxidative capacity, we supposed that BAT oxidative metabolism could be altered. This prompted us at firstly analyzing BAT mitochondria at ultrastructural level through transmission electron microscopy. In WT mice typical mitochondria (abundant, large, and rich in cristae) were present in BAT, while in KIKO mice the mitochondria appeared lower in number and enlarged with disorganized and thickened cristae (**Fig. 3A**). We then compared the basal oxygen consumption in mitochondria isolated from BAT and as expected the oxygen consumption was lower in KIKO than WT mice (**Fig. 3B)**.

**Fig. 3.**
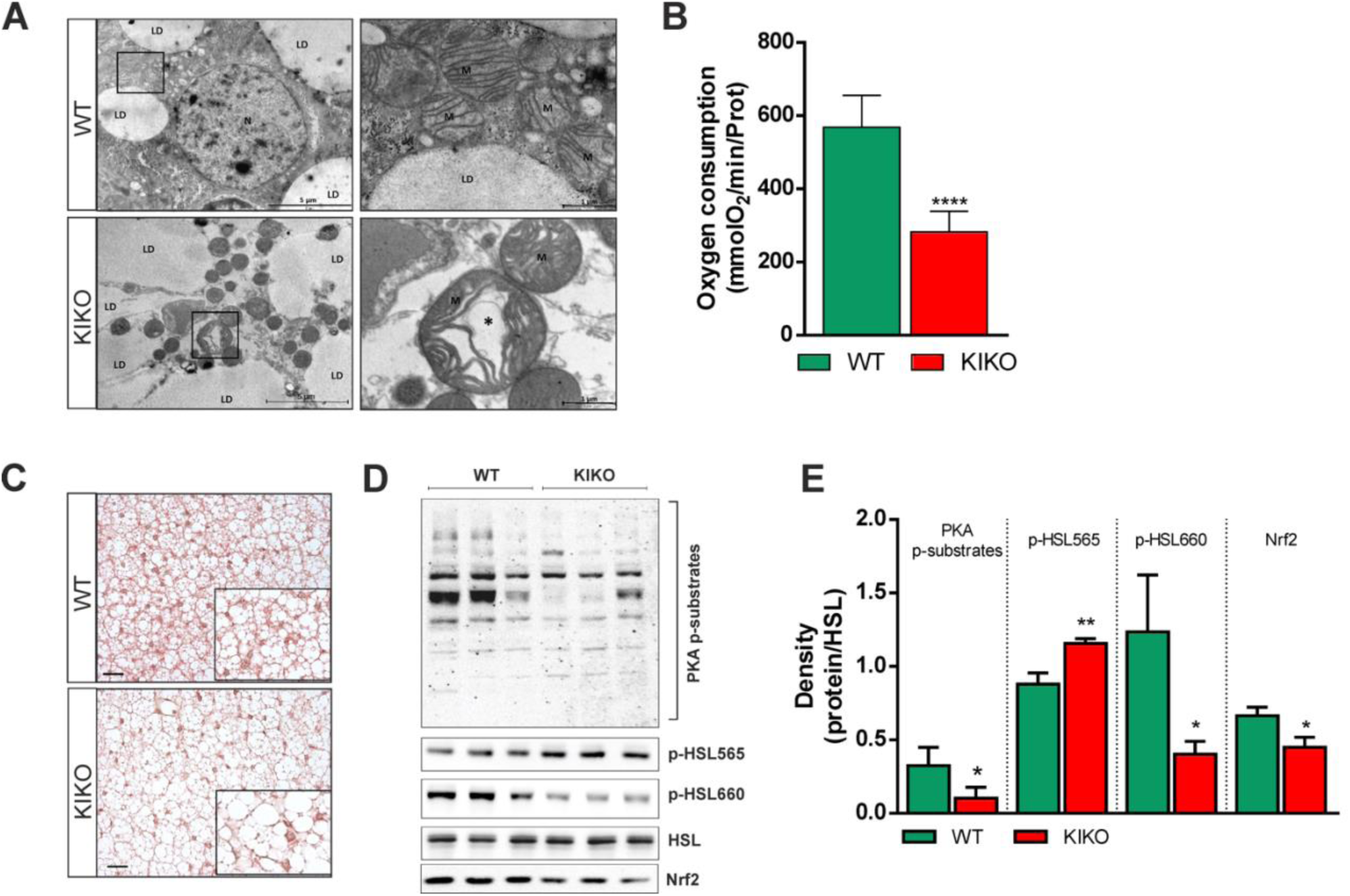
Mitochondria of KIKO mice show altered mitochondrial morphology and respiratory function. **(A)** Representative transmission electron microscopy pictures of BAT from 6-months old WT and KIKO mice are reported in *left panels* (5800x). Images with high power field (18500x) are reported in *right panels*. A representative mitochondrion of KIKO mice with altered ultrastructure was evidenced with an asterisks (*). LD, lipid droplets; N, nucleus; M, mitochondria. **(B)** Oxygen consumption determined through a polarographic method on crude mitochondria isolated from BAT of 6-months old mice (n=6 each group; ****p<0.0001 vs WT). **(C)** Representative BAT histology images after staining with H&E (n=3 each group). Scale bars, 25μm. (**D, E**) Representative immunoblots (D) of total HSL, phospho-active (p-HSL660) and phospho-inactive HSL (p-HSL565), PKA phosphorylated substrates (PKA p-substrates) and Nrf2 in total BAT homogenates, and densitometric analyses of the immunoreactive bands (E). HSL was used as loading control (n=6 each group, *p<0.05, **p<0.01 vs WT).

Accumulation of lipid droplets (LDs) and an increased lipogenesis have been previously described in fibroblasts of FRDA patients and cardiomyocytes from KIKO mice (Coppola et al., 2009). In line with these data, we found that BAT of KIKO mice has higher lipid droplets dimension with respect to WT mice (**Fig. 3C**). We have tested whether this event was dependent on defective lipolysis that importantly contributes to LDs turnover. BAT lipolysis strongly relies on cAMP-dependent protein kinase A (PKA) that coordinates lipolytic machinery activation through hormone/phospho-dependent mechanisms. To this end, we evaluated the level of the PKA phosphorylated substrates as well as the lipases directly responsible for LDs degradation. As reported in the immunoblot in **Fig. 3D**, PKA substrates were reduced in their phosphorylated levels, suggesting a general inhibition of PKA activity. Among the PKA substrates the hormone-sensitive lipase (HSL) is included, as it is activated via the phosphorylation of Ser660. While the basal levels of HSL remained unaltered, the phospho-active levels of HSL were lower in KIKO compared to WT mice (**Fig. 3D, E**). The levels of phosphorylation at Ser565 (phospho-inactive form) were instead increased, indicating an impaired basal HSL activity. According to what reported in other studies on mouse model and patients (Shan et al., 2013;Sahdeo et al., 2014), we also found a significant decrease of the transcription factor Nrf2 (**Fig. 3D, E**) that, besides being an up-stream modulator of the expression of antioxidant genes and protecting against oxidative stress, positively regulates enzymes involved in mitochondrial fatty acids oxidation (Esteras et al., 2016).

### KIKO mice display affected expression of genes related to lipid utilization and thermogenesis

Considering the alteration of mitochondrial respiration and lipid catabolism observed in BAT, we hypothesized a compromised thermoregulatory activity in KIKO mice. To test this, we next exposed mice to cool temperature (4°C) for 12 h and a lower body temperature was measured in KIKO mice both at thermoneutrality as well at 4°C (**Fig. 4A)**. Remarkably, BAT of cold-exposed mice showed a that KIKO mice maintained a higher LD size with respect to WT mice (**Fig. 4B**). In particular, the measurement of LD size at RT confirmed their higher dimension in KIKO mice compared to WT mice (**Fig. 4C**). Cold exposure promoted a marked reduction of LDs diameter both in WT and KIKO mice; however, in cold-exposed KIKO mice LDs were larger than WT mice (**Fig. 4C**).

**Fig. 4.**
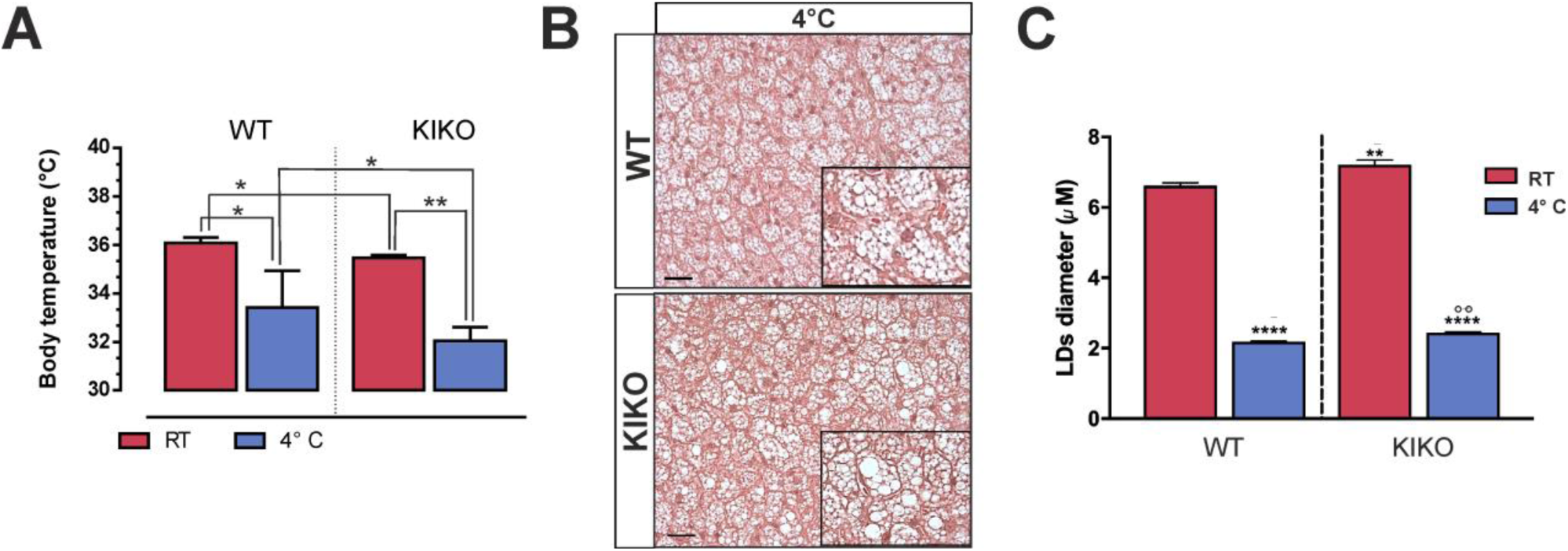
KIKO mice have reduced cold tolerance. **(A)** Body temperature measured in 6-months old mice after 12 h cold exposure (n=6 each group;*p<0.05, **p<0.01). **(B)** Representative histology images of BAT from mice exposed to cold after staining with H&E (n=3 each group). Scale bars, 25 μm. **(C)** Lipid droplets (LDs) diameter measurement in three fields for each BAT histology through ImageJ (300 droplets total at least for each sample). BAT of WT and KIKO mice at RT (see Fig. 4C) or exposed to cold (B) for 12 h were analyzed (n=3, ****p<0.0001 vs RT, **p<0.001 vs WT mice at RT, °°p<0.001 vs WT mice at 4°C).

To more deeply decipher the cause(s) of cold intolerance, we performed BAT transcriptome profiling using an ultra-deep unbiased RNA sequencing (RNA-seq) approach. The RNA samples analyzed were prepared from BAT of WT and KIKO mice maintained at RT or exposed to cold for 12 hrs. A summary of the results obtained through RNAseq is reported in the volcano plots depicted in **Fig. 5A**. Through pair-wise differential gene expression, we found that about 200 genes were up-regulated (Log2FC>0.58; p<0.05) upon cold exposure both in WT and KIKO mice. In order to determine which biological processes or pathways were overexpressed, the list of up-regulated genes was used as the input for a functional enrichment analysis. By using the plugin ClueGo in the Cytoskape 3.7.1 platform, as expected we found that the response to temperature stimulus was the biological processes significantly up-regulated either in WT or KIKO mice exposed to cold with respect to their controls, even though KIKO mice showed a lower enrichment with respect to WT mice (**Fig. 5B**). The biological processes found overrepresented in WT mice were positive regulation of cold induced thermogenesis, triglyceride metabolic process (including lipolytic genes), monocarboxylic acid transport (including fatty acid transporters) and mitochondrial gene expression (**Fig. 5B, left panel**), in line with the notion that fatty acids transport and degradation, and expression of mitochondrial proteins accompany thermogenesis. Interestingly, genes mapping to categories pertaining purine nucleotide and monocarboxylic acid biosynthetic processes (including fatty acids biosynthesis genes) were found overexpressed in KIKO mice exposed to cold compared to KIKO mice at RT (**Fig. 5B, right panel**). Moreover, in cold-exposed KIKO mice we found an enrichment of reactive oxygen species (ROS) biosynthetic process and cellular response to xenobiotic stimulus (**Fig. 5B, right panel**).

**Fig. 5.**
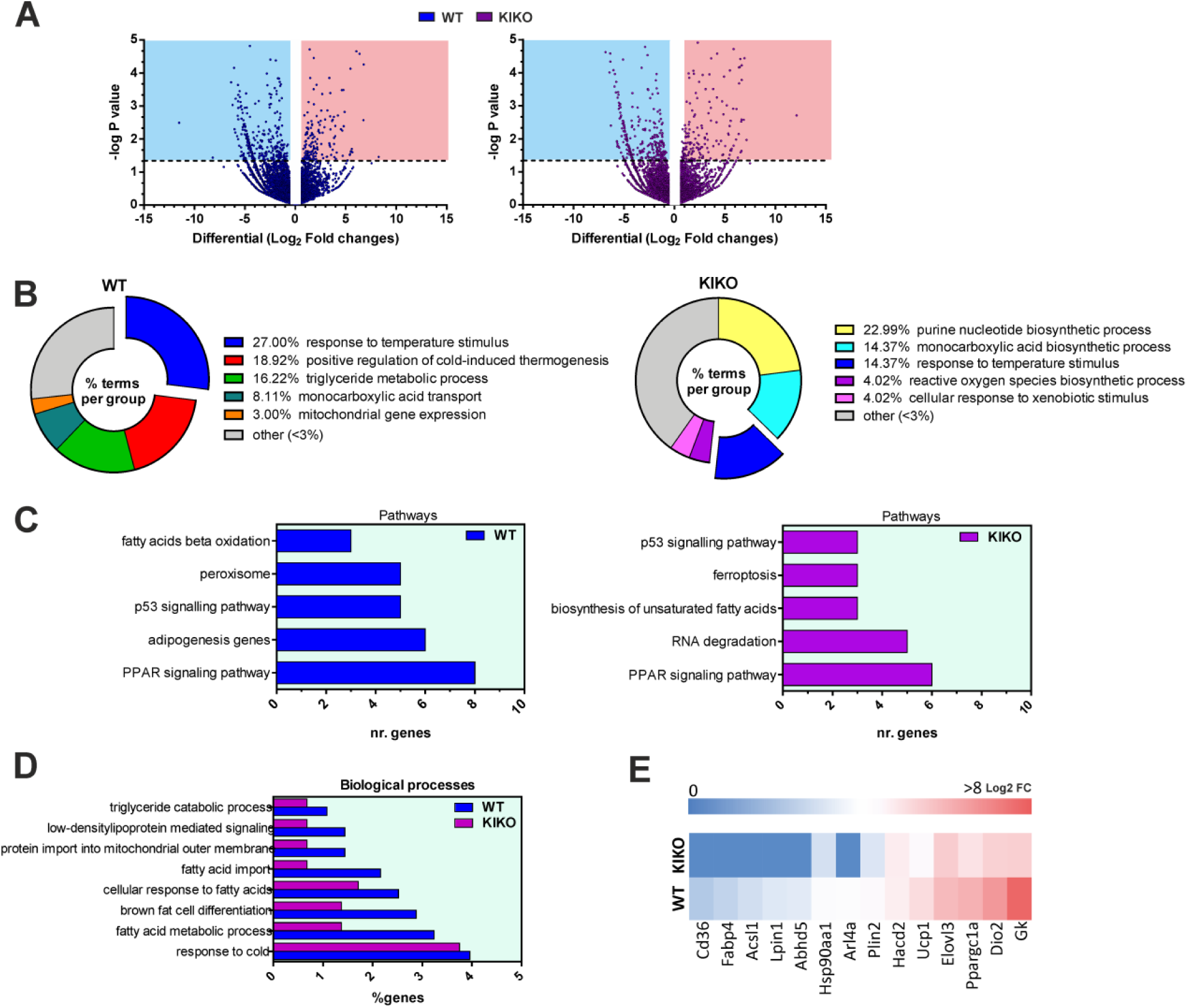
RNAseq analysis revealed reduced expression of genes related to BAT activity in KIKO mice. (**A**) Volcano plot representation of differential gene expression analysis in BAT of 6-months old WT (*upper panel*) and KIKO mice (*lower panel*) exposed to cold for 12 h. Light blue and red squares mark the genes with significantly decreased or increased expression respectively (p<0.05, n=3). (**B, C**) Biological processes (B), KEGG and Wiki pathways (C) significantly up-regulated in BAT of WT (*left panels*) and KIKO mice (*right panels*) upon cold exposure determined through ClueGo plugin of Cytoscape v3.7 platform (Benjamini-corrected p values < 0.05). **(D)** Comparative analysis of significantly up-regulated biological processes (p<0.05) in WT mice and KIKO mice determined through Funrich v3.0. **(E)** Heatmap showing representative genes significantly up-regulated (p<0.05) in WT mice compared to KIKO mice.

The analysis of KEGG and Wiki pathways showed that genes belonging to PPAR signaling, notably induced during thermogenesis, were significantly overrepresented both in cold-exposed WT and KIKO mice; however, the number of genes found in this pathway was lower in KIKO than WT mice (**Fig. 5C**). The same trend was observed for p53 signaling pathway. Among the top enriched pathways, WT mice also had adipogenesis, peroxisome and beta-oxidation (**Fig. 5C, left panel**); by contrast, RNA degradation, biosynthesis of fatty acids and ferroptosis were among the enriched pathways found in BAT of cold-exposed KIKO mice (**Fig. 5C, right panel**). Ferroptosis is a more recently recognized cell death caused by iron overload (Dixon et al., 2012;Stockwell et al., 2017), suggesting that in KIKO mice the thermogenic response of BAT is blunted and likely accompanied by iron/ROS-induced cell death.

To complement our analysis, we alternatively examined transcriptomic data through Funrich v3.0 and found that the response to cold was the biological process significantly and similarly enriched in WT and KIKO mice upon cold exposure (**Fig. 5D**). Brown fat cell differentiation and other biological processes related to in lipid metabolism were found overrepresented both in WT and KIKO mice. However, in KIKO mice a lower percentage of genes pertaining to these processes was found compared to WT mice (**Fig. 5D**). In **Fig. 5E**, a heatmap including some representative genes strictly involved in adipocyte differentiation, heat production and thermogenic-related metabolic re-adaptation is illustrated. The heatmap shows that KIKO mice undergo a lower up-regulation of these genes compared to WT mice, arguing that cold intolerance of KIKO mice could be ascribed to defective activation of adipocytes differentiation, thermogenic pathway, impaired BAT lipid catabolism and/or oxidative capacity.

Transcriptomic results were confirmed by performing qPCR analyses. In particular, we confirmed the downregulation of FXN mRNA in KIKO mice and found a significant downregulation of the expression of other mitochondrial genes such as Tfam, Mtco1, Nd1, Atp6 and Nd4 (**Fig. 6A**). Upon cold exposure, WT mice showed up-regulation of Ucp1, Tfam, Nd1 and Atp6. In KIKO mice an increase of Ucp1, Tfam, Nd1 and Atp6 was also achieved; however, the level of upregulation was lower than WT mice.

**Fig. 6.**
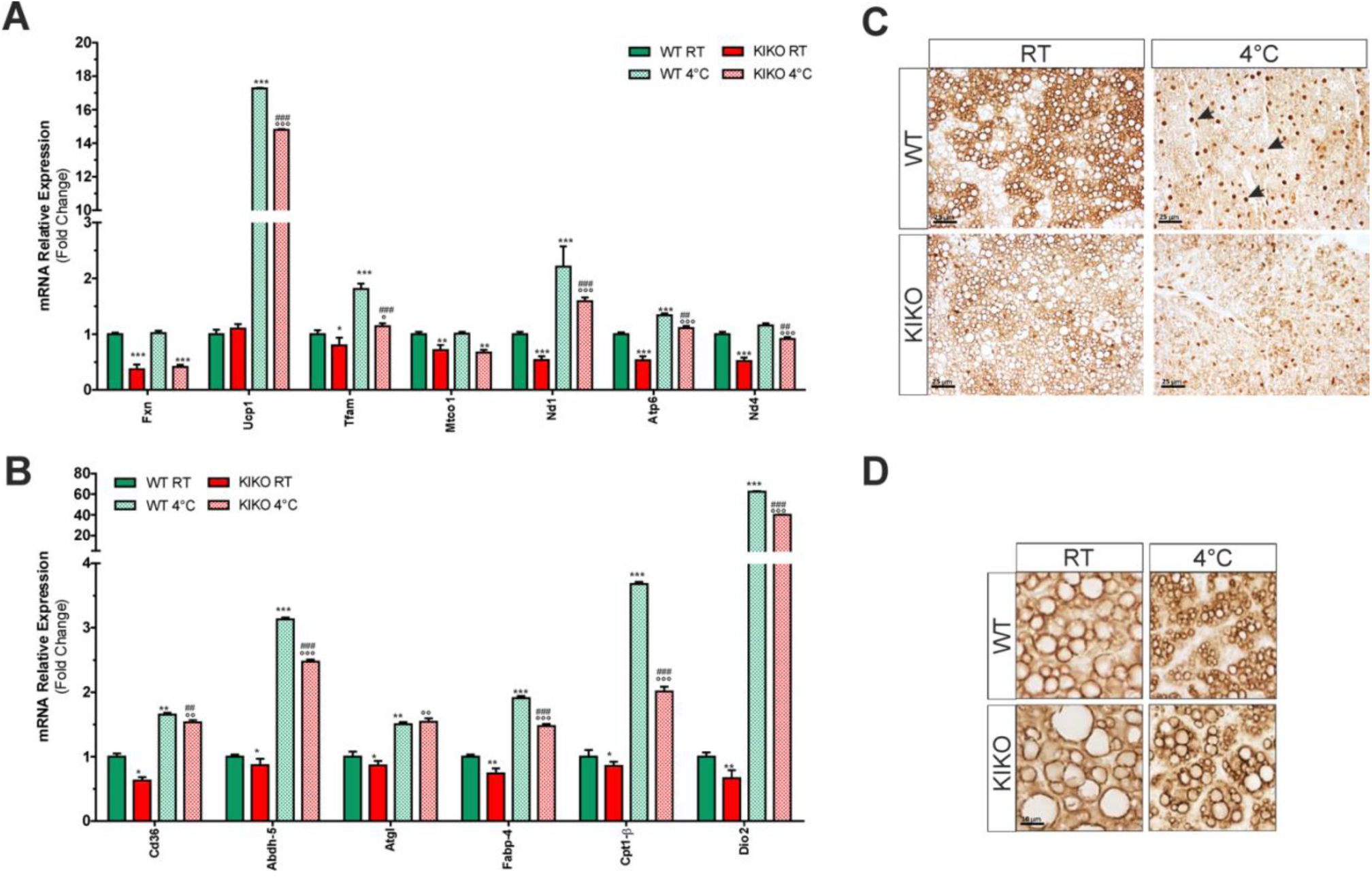
The expression of genes related to mitochondrial respiration, thermogenesis and lipolytic pathways are altered in BAT of KIKO mice. (**A, B**) RT-qPCR analysis of mitochondrial genes (A) and genes related to lipid metabolism (B) in BAT of 6-months old mice (n=6; *p<0.05, **p<0.01, ***p<0.001 vs WT; °p<0.05,°°p<0.01,°°°p<0.001 vs KIKO; ^##^p<0.01, ^###^p<0.001 vs WT at 4°C). (**C, D**) Representative immunohistochemical analyses of phospho-PKA substrates (C) and perilipin-1 (D) (n=3 mice each group). Black arrows (C) indicate some nuclei showing positivity to the immunostaining of phospho-PKA substrates.

We also confirmed the alterations in the expression level of genes related to lipid metabolism and thermogenesis in KIKO mice. Actually, as showed in **Fig. 6B**, the expression of genes implicated in fatty acids transport (i.e. Cd36, Fabp4, Cpt1b) and thermogenesis (i.e. Dio2) resulted downregulated in KIKO mice. A slight but significant decrease of the lipase responsible for the first step of triglyceride lipolysis Atgl and its enhancer Abdh5 was also found, indicating an alteration of the triglyceride catabolism and lipid signaling. Upon cold exposure, overall these genes resulted up-regulated both in WT and KIKO mice; however, in KIKO mice Cd36, Abdh5, Fabp4, Cpt1b and Dio2 resulted up-regulated at minor extent (**Fig. 6B**).

Through immunohistochemical analyses we confirmed the alteration of PKA lipolytic pathway in BAT of KIKO mice. Indeed, a decrease of PKA kinase activity was occurring already at RT and a further reduction of phospho-PKA substrates was found at 4°C with respect to WT mice (**Fig. 6C**). In particular, at RT the BAT of WT mice shows brown adipocytes with a diffused and stronger staining with respect to BAT of KIKO mice. Upon cold exposure, the immunostaining of phospho-PKA substrates was predominantly nuclear, in line with the notion that PKA can migrate into the nucleus during thermogenesis (Rim et al., 2004) and phosphorylate a number of transcriptional regulators (e.g. Ppargc1a) and transcription factors triggering adipogenesis and thermogenesis as well as mitochondrial oxidative activity and biogenesis (e.g. CREB) (Cannon and Nedergaard, 2004;Baldelli et al., 2014). Notably, the nuclear staining of phospho-PKA substrates was markedly reduced in BAT of cold-exposed KIKO mice (**Fig. 6C**). Moreover, in these mice we found an increase of LD-associated protein perilipin-1 (**Fig. 6D**), which is an inhibitor of triglyceride degradation by HSL (Grahn et al., 2013), thus highlighting the dysfunction of lipolysis in BAT of KIKO mice.

### FXN deficiency leads to impaired thermogenesis and adipogenesis in cultured adipocytes

To confirm that FXN deficiency affects lipid utilization and thermogenesis, we moved to an *in vitro* system. In cultured T37i brown adipocytes, we downregulated FXN levels through RNA interference by transfecting a pool of siRNA against FXN mRNA (FXN-). Transfection of T37i cells with a pool of scramble siRNAs were used as control (Scr). To induce thermogenic cascade, T37i were treated with isobutyl methylxanthine (IBMX), a nonspecific inhibitor of phosphodiesterase (PDE), which enhances the intracellular cAMP levels, inducing a constitutive activation of PKA. As reported in **Fig. 7A**, the analysis of LDs evidenced that lipolysis was likely blunted as consequence of FXN deficiency as LDs dimension was larger in FXN-cells than Scr cells both in untreated and IBMX-treated cells. **Fig. 7B** shows that FXN downregulation did lead to the inhibition of lipolysis, as reduced levels of phospho-active HSL (p-HSL660), a known target of PKA activity, was elicited both upon basal condition and IBMX treatment. In line with the transcriptomic data obtained in BAT of KIKO mice, PPARγ, a thermogenic transcription factor regulating mitochondrial biogenesis and lipid catabolism, was reduced in FXN-cells under resting conditions and, in contrast to Scr cells, it remained down-regulated upon IBMX treatment (**Fig. 7C**). Importantly, UCP1 protein was efficiently induced by IBMX in Scr cells but not in FXN-cells. Mitochondrial proteins including VDAC and TOM20 were also reduced in FXN-cells both prior and after IBMX treatment (**Fig. 7C**). RT-qPCR analyses confirmed the impairment of thermogenic program, as the mRNA expression of thermogenic markers including Cox7a, Ppargc1a, Cidea and Cd36 were significantly reduced in FXN-cells both under basal condition and upon IBMX treatment with respect to Scr cells (**Fig. 7D**). In line with these data, reduced oxygen consumption was also recorded in FXN-cells (**Fig. 7E**). Since RNAseq analyses pointed to a defective induction of adipogenic genes upon cold exposure, we then attempted at evaluating whether adipocyte differentiation potential could be impaired in KIKO mice. We hence isolated stromal vascular cells (SVCs) from subcutaneous adipose tissue, which is highly enriched with primary adipocyte precursors. Upon certain circumstances (i.e. rosiglitazone treatment), SVCs may differentiate in brown-like adipocytes displaying high mitochondrial mass and expressing detectable levels of Ucp1 (Aune et al., 2013). As reported in **Fig. 8A**, WT SVCs were efficiently differentiated in adipocytes as a great number of cells show adipocyte morphology with an evident accumulation of LDs. On the contrary, SVCs from KIKO mice show affected adipogenic capacity as a lower number of adipocytes was obtained. As expected, RT-qPCR analyses evidenced a marked reduction of FXN mRNA levels in KIKO with respect to WT adipocytes that was accompanied by a significant down-regulation of genes related to adipogenesis, thermogenesis and mitochondrial biogenesis including Pparγ, Ppargc1a, Ucp1, Cd36 and Mtco1 (**Fig. 8B**).

**Fig. 7.**
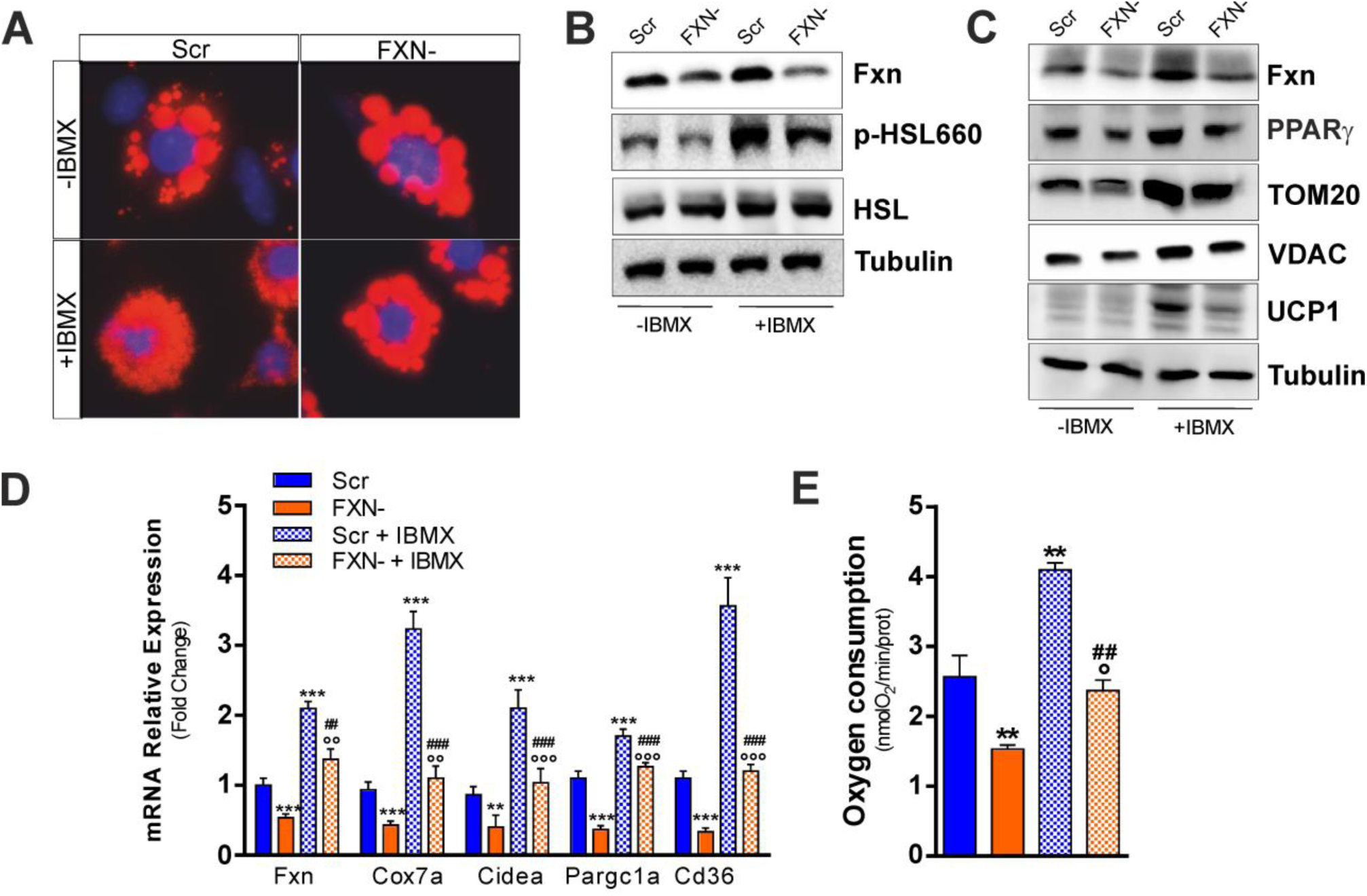
FXN deficiency impairs thermogenic program also in a brown adipocytes cell line. (**A**) Representative immunofluorescence analysis of lipid droplets after staining with the neutral lipid probe Nile Red in T37i brown adipocytes transfected with a pool of siRNAs targeting FXN mRNA (FXN-) or with a pool of Scr siRNAs (Scr) (n=4). (**B, C**) Representative immunoblots of FXN, total and phospho-active HSL (p-HSL660) (B), PPARγ, TOM20, VDAC and UCP1 (C) (n=4). **(D)** RT-qPCR analysis of genes implicated in thermogenesis (n=4; **p<0.01,***p<0.001 vs Scr; °°p<0.01, °°°p<0.001 vs FXN-; ^##^p<0.01, ^###^p<0.001 vs Scr + IBMX). **(E)** Cellular oxygen consumption determined through a polarographic method (n=4, **p<0.01 vs Scr, °p<0.05 vs FXN-; ^###^p<0.001 vs Scr + IBMX).

**Fig. 8.**
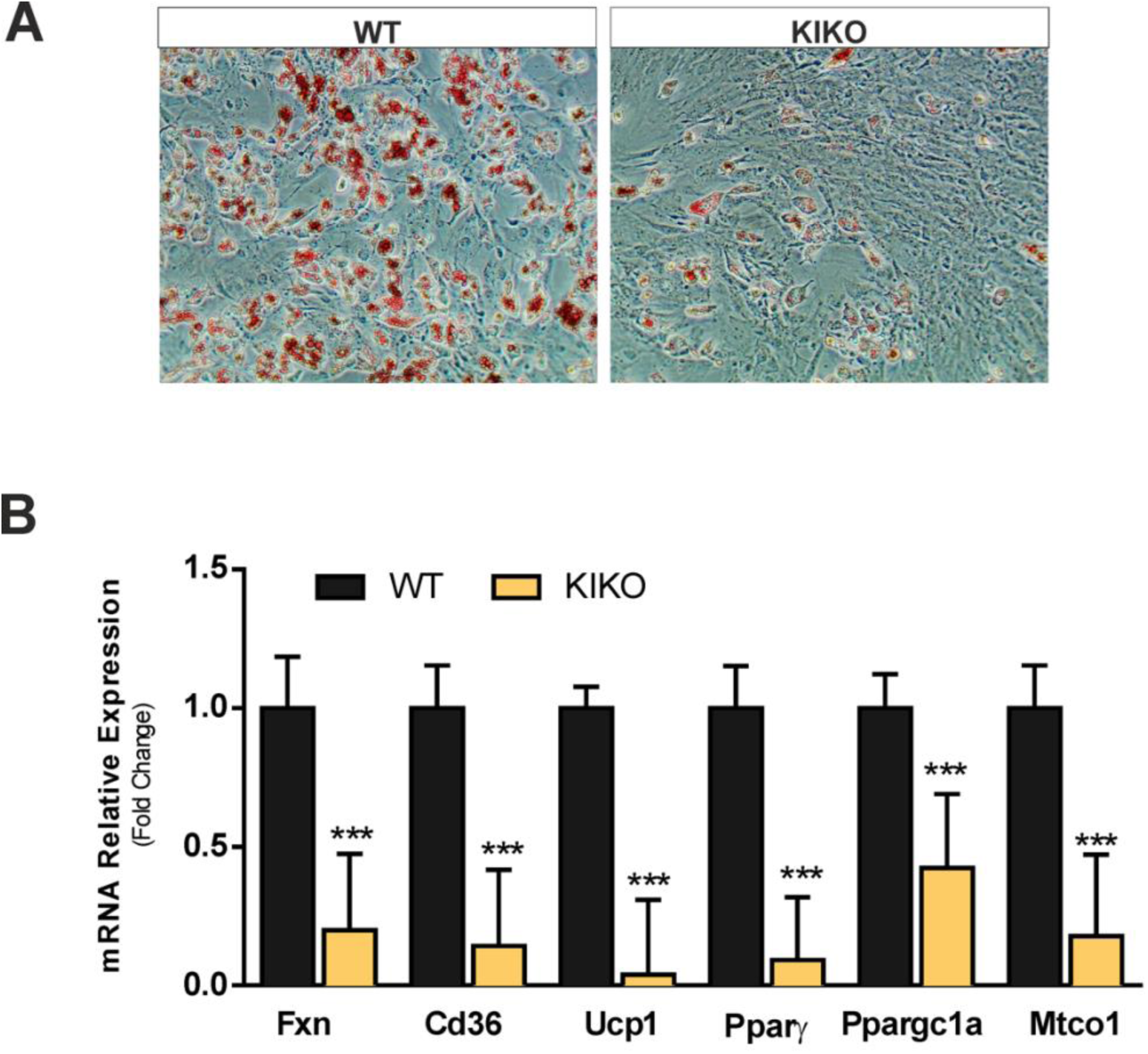
KIKO mice show affected adipogenic potential. **(A)** Representative images of stromal vascular cells (SVCs) obtained from mouse adipose depots of WT or KIKO mice, differentiated in adipocytes and stained with Oil Red O to detect lipid droplet-containing mature adipocytes (n = 3). **(B)** RT-qPCR analysis of genes implicated in adipogenesis and thermogenesis in SVCs differentiated in adipocytes (n = 6; ***p<0.001 vs WT).

## DISCUSSION

Abnormal glycemic control, increased cholesterol and triglycerides levels as well as type 2 diabetes are more frequent in FRDA patients than in the general population and concur in the severity of FRDA (extensively reviewed in (Tamarit et al., 2016)). In this work we demonstrated KIKO mice show several diabetes-related hallmarks, including hyperlipidaemia, altered tolerance to glucose, and elevated circulating level of leptin, a peptide hormone secreted by adipose tissue whose increase is strictly related to insulin resistance and low-grade inflammatory states (Francisco et al., 2018). Overall these metabolic changes make this model suitable to decipher the events contributing to disease severity and find novel druggable target to overcome type 2 diabetes development in FRDA patients.

In the last decade, the scientific community has put a spotlight on BAT as a crucial player in the control of energy metabolism, being the main glucose and lipid clearance organ with the highest mitochondrial fatty oxidation rate (Doh et al., 2005;Bartelt et al., 2011). We have reported that BAT mitochondria of KIKO mice show a significant ultrastructural alteration, lower abundance and oxygen consumption and decreased mRNA expression of electron transport chain complex subunits compared to WT mice. Upon cold exposure KIKO mice show affected adaption to cold with an overall less efficient up-regulation of canonical thermogenesis-related genes and pathways including mitochondrial and lipid metabolism genes and PPAR signaling pathway.

It has been reported that FRDA patients show LDs accumulation in fibroblasts (Coppola et al., 2009). LDs accumulation was observed also in FXN deficient cultured cardiomyocytes (Obis et al., 2014) and heart of FRDA mouse models (Puccio et al., 2001;Stram et al., 2017). Hepatic steatosis in mice with a liver-specific FXN ablation (Martelli et al., 2012) and altered lipid metabolism associated with increased LDs in glial cells of the drosophila FRDA model (Navarro et al., 2010) were also observed. Down-regulation of PPARγ/PGC-1α pathway and up-regulation of lipogenic genes have been previously proposed among the mechanisms leading to LDs accumulation (Sutak et al., 2008;Coppola et al., 2009). We have evidenced that FXN deficiency leads to LDs accumulation in BAT as well. Notably, the size of LDs is the result of the balance between lipid demolition and deposition. Since BAT requires plentiful FAs, which are its primary substrate, lipolysis is activated during BAT thermogenesis (Cannon and Nedergaard, 2004;Nedergaard et al., 2011;Townsend and Tseng, 2014). We have demonstrated that impaired lipid degradation could contribute to LDs accumulation, as PKA-dependent lipolysis resulted significantly affected both under resting condition and upon cold exposure. As consequence, a significant reduction of the active form of HSL and increase of the LD-associated inhibitor of lipolysis perilipin was observed in BAT of KIKO mice.

Importantly, besides acting as fuels to sustain the electron transport chain flow, fatty acids are involved in the regulation of UCP1, or directly activate UCP1-mediated energy dissipation and heat production (Cannon and Nedergaard, 2004). The impaired lipid degradation observed in KIKO mice likely limits the funneling of fatty acids into mitochondria with consequent affected capacity to produce heat. Accordingly, fatty acids beta-oxidation was not disclosed among the enriched pathways in BAT of cold-exposed KIKO mice and a diminished expression of the mitochondrial fatty acid carrier Cpt1b was found. Notably, intracellular purine nucleotides exert an inhibitory action on UCP1 protein (Fromme et al., 2018). It was recently demonstrated that upon thermogenic stimuli, brown adipocyte expression of enzymes implicated in purine metabolic remodeling is altered (Fromme et al., 2018). In particular, an overexpression of genes implicated in purine nucleotide degradation was found that was associated with a down-regulation of genes involved in purine nucleotide synthesis. In BAT of KIKO mice exposed to cold we found an overrepresentation of genes pertaining purine nucleotide biosynthetic process. Therefore, these data collectively indicate that UCP1, in addition to being up-regulated at lesser extent than WT mice, is less efficient in impinging heat production. To exclude that the alterations found in BAT of KIKO mice could be the results of systemic adaptive responses to FXN deficiency, we validated our findings in cultured brown adipocytes downregulating FXN. By igniting thermogenic program via the PKA agonist IBMX we confirmed that FXN deficiency leads to the inhibition of lipolysis and affected up-regulation of genes related to the thermogenic cascade.

Another process having a critical role in maintaining thermogenic efficiency of BAT is the differentiation of resident brown fat adipocytes *de novo* that assures BAT regeneration and avoids its loss (Birerdinc et al., 2013). Transcriptomic data evidenced brown fat cell differentiation and adipogenesis as biological process and pathway affected upon cold exposure in BAT of our mouse model. We confirmed such hypothesis by analyzing the adipogenic potential of stromal vascular cells isolated from mouse adipose depots that resulted unable to fully differentiate in thermogenic adipocytes in KIKO mice.

It is important to notice that a less proficient up-regulation of p53 signaling pathway in KIKO mice was observed in BAT upon cold exposure. p53 is a stress responsive transcription factor that was reported to exert a positive regulatory effect on brown adipocyte differentiation (Al-Massadi et al., 2016) and BAT thermogenesis (Molchadsky et al., 2013). This evidence corroborates our idea that BAT dysfunction could at least in part depend on altered capacity of FXN deficient adipocyte precursors to efficiently differentiate and express mitochondrial and thermogenic genes.

Even though not still adequately investigated in the present work, upon cold stress condition the KEGG pathway of ferroptosis was significantly up-regulated in KIKO mice along with genes related to ROS metabolism. This is not surprising as we found a decrease of the Nrf2 protein, and oxidative stress, iron overload and ferroptosis have been claimed among the main pathogenic factors in FRDA (Shan et al., 2013;Petrillo et al., 2017;Abeti et al., 2018;Cotticelli et al., 2019). Interestingly, Nrf2, besides being an up-stream modulator of the expression of antioxidant genes and protection against oxidative stress/ferroptosis (Faraonio et al., 2006;Dodson et al., 2019), positively regulates enzymes involved in mitochondrial fatty acids oxidation (Esteras et al., 2016) and adipogenesis (Pi et al., 2010;Hou et al., 2012). Based on this, Nrf2 inducers, already proposed as FRDA therapeutics (Petrillo et al., 2017;Abeti et al., 2018) could be also advantageous for preserving BAT integrity and mitigating metabolic disturbances of FRDA patients via a BAT-dependent manner. Accordingly, Nrf2 targeting has been proposed as promising for treating type 2 diabetes (David et al., 2017).

BAT studies carried out in human fetuses and infants indicate that the tissue is widely distributed during these developmental stages and that the thermogenic capacity of BAT develops with gestational age reaching its maximum in infancy and early childhood when the demands for thermogenesis can be expected to be especially high (Lidell, 2019). Several studies suggest that BAT dysfunction during gestation and early childhood negatively influences metabolic health predisposing to type 2 diabetes development later in life (Liang et al., 2016;Entringer et al., 2017;Lettieri-Barbato et al., 2017). Therefore, it is possible to postulate that FRDA patients would have experienced BAT dysfunction early in life leading to disruption of systemic metabolic homeostasis.

In conclusion, by deeply characterizing KIKO mice at metabolic level we have provided multiple lines of evidence that FXN deficiency in mice leads to clinical-pathological features parallel to those observed in diabetic patients. Among the metabolic parameters we have evidenced that the lipolytic and thermogenic activities of BAT are reduced, thus providing the possibilities of targeting BAT that might result in therapeutic benefits in FRDA.

## MATERIALS AND METHODS

### Animals

Mouse experimentation was conducted in accordance with accepted standard of humane animal care after the approval by relevant local (Institutional Animal Care and Use Committee, Tor Vergata University) and national (Ministry of Health, license n°324/2018-PR) committees. Unless otherwise stated, mice were maintained at 21.0 ± °C and 55.0 ± 5.0% relative humidity under a 12 h/12 h light/dark cycle (lights on at 6:00 AM, lights off at 6:00 PM). Food and water were given *ad libitum*. Experiments were carried out according to institutional safety procedures.

Knock-in knock-out (KIKO) mice were purchased from Jackson Laboratories (#012329). Littermate C57BL/6 mice (WT) were used as controls. Researchers were blinded to genotypes at the time of testing. Only WT and KIKO male mice were used and divided in the following groups: 1) 6 months-old WT and KIKO mice (n=12 each group); 2) 8 months-old WT and KIKO mice (n=12 each group); For cold exposure experiments additional male WT and KIKO at 6-months of age were divided in the following groups: 1) WT and KIKO mice maintained at room temperature (n=6 each group); 2) WT and KIKO mice maintained at 4°C for 12 h (n=6 each group). Rectal temperature was measured by high precision (± 0.1 °C) rectal probe for mice (RET-3, ThermoWorks, Alpine, UT, USA).

### Bio-clinical and adipokine analyses

Prior to bio-clinical analyses, mice were starved for 2 h. After blood collection, bio-clinical analyses were performed by colorimetric methods. In particular, glucose, cholesterol, triglycerides, creatinine, total plasma proteins, albumin and urea were measured through the automatized KeyLab analyzer (BPCBioSed, Italy) using specific assay kits (BPCBioSed).

Serum adiponectin, FGF21 and leptin levels were measured through a Mouse Magnetic Luminex Screening Assay (R&D System, Minneapolis, MN, USA).

For the glucose tolerance test (OGTT), mice were subjected to fasting for 12 h, followed by oral gavage with 2 g of dextrose/kg body mass. At the indicated time points, blood was collected from the tail vein and glycaemia measured using a glucometer (Bayer Countur XT, Bayer Leverkusen, Germany).

### Indirect calorimetry

Indirect calorimetry performed using LabMaster (TSE Systems, Bad Homburg, Germany) as previously described (Lettieri-Barbato et al., 2018). Oxygen consumption (VO2), carbon dioxide production (VCO2) and Resting energy expenditure (REE) were recorded every 15 min for 24h, and the data were averaged for each mouse.

### Histochemical analysis

Formalin-fixed paraffin-embedded (BAT explants were cut in 3 μm sections and stained with hematoxylin and eosin (H&E) prior microscope analysis. Lipid droplet diameters were measured in three fields (300 droplets total at least) with ImageJ. For correlative purpose, an average score was derived for each sample. Perilipin and phospho-PKA substrate levels were investigated by immunohistochemistry on tissue sections. After antigen retrieval with Citrate Buffer (pH 6.0), sections were incubated at room temperature with the following primary antibodies: rabbit polyclonal antibody anti-Perilipin diluted 1:1000 (9349T, Cell Signalling Technology) and rabbit polyclonal antibody anti-phospho-PKA substrates diluted 1:100 (9621S, Cell Signalling Technology). Negative controls were obtained by omitting primary antibodies. Immunohistochemical reactions were visualized by DAB as the chromogen from MACH 1 Universal HRP-Polymer Detection (Biocare Medical, Concord, MA, USA).

### Ultrastructural Analysis

BAT samples from animal models were fixed in 2.5% glutaraldehyde in 0.1 M cacodylate buffer for the morphological study. After fixation, dehydration and impregnation, samples were included in epoxy resins and acrylic, cut at the ultramicrotome and processed for the ultrastructural study of brown adipocytes by electron microscopy. In particular, morphological aspects of the mitochondria, such as the presence and density of cristae, were evaluated. Ultrastructural images were collected with a Transmission Electron Microscope FEI TecnaiTM (Hillsboro, Oregon, USA), equipped with a dedicated Imaging Software System.

### Cell and mitochondria oxygen consumption

T37i cell line was gently provided by Prof. Marc Lombes (Inserm U693, Paris, France), cultured and differentiated as described by Nakae et al., 2008 (Nakae et al., 2008). Differentiated T37i brown adipocytes were transfected with a pool of 3 target-specific siRNAs against FXN mRNA FXN or scramble siRNAs (Santa Cruz Biotechnology, Dallas, TX, USA) by using DeliverX Plus kit (Affymetrix, Santa Clara, CA, USA). Treatment with isobutyl methyl xanthine (IBMX) was carried out at concentration of 0.5 mM for 4 h. For lipid droplet detection, cells were stained with Nile Red (0.25 μg/ml,10 min). Staining with Hoechst 33342 (1 μg/ml, 10 min) was used to counterstain nuclei. Images were visualized by Nikon Eclipse TE200 epifluorescence microscope (Nikon, Florence, Italy) connected to a CCD camera. Images were captured under constant exposure time, gain and offset. Stromal vascular cells (SVCs) were isolated from subcutaneous adipose tissue and differentiated in brown-like adipocytes according to Aune et al (Aune et al., 2013). Oil Red O was used to detect intracellular triglycerides content as previously described (Lettieri-Barbato et al., 2018).

Oxygen consumption was determined in cells and crude mitochondria and by using the Oxygraph Plus oxygen electrode system (Hansatech Instruments Ltd., Norfolk, UK). Intact cells were resuspended (1 × 10^6^/ml) in culture medium without FBS. Crude mitochondria were isolated from BAT as previously described (Lettieri Barbato et al., 2015) and resuspended in an appropriate mitochondrial activity buffer (70 mM sucrose, 220 mM mannitol, 2 mM HEPES buffer, 5 mM magnesium chloride, 5 mM potassium phosphate, 1 mM EDTA, 5mM succinic acid, and 0.1% fatty acid free bovine serum albumin, pH 7.4). Oxygen consumption rate was recorded at 37° C for 10 min and normalized for protein concentration.

### RNA-seq data expression quantification and functional enrichment analysis

The quality of the single-end reads was evaluated with *FastQC* v.0.11.5 (https://www.bioinformatics.babraham.ac.uk/projects/fastqc/). All the *fastqc* files were filtered to remove low quality reads and adapters with *Trimmomatic* v.0.36 (Bolger et al., 2014). The resulting reads were mapped to the *Mus musculus* genome (GRCm38) with *HISAT2* v.2.1.0 (Kim et al., 2015) using default parameters, while *Stringtie* v1.3.4d (Pertea et al., 2015) was applied to the BAM files obtained with *HISAT2* to generate expression estimates and to quantify the transcript abundance as transcripts per kilobase per million of mapped reads (TPM). The count matrices generated by *Stringtie* were imported in *R* where differential expression analysis was performed using the *Deseq2* package (Love et al., 2014) to compare the two different conditions. The functional annotation was performed through the *AnnotationDbi* R library (http://bioconductor.org/packages/release/bioc/html/AnnotationDbi.html).

Differential expressed genes were selected with threshold of Log2FC>0.58 (p<0.05). Functional enrichment analysis including GO and Kyoto Encyclopedia of Genes and Genomes (KEGG) pathway was performed by using the ClueGo plugin of the Cytoscape v3.7.1. ClueGo settings were: enrichment only, Benjamini-Hochberg false discovery rate (FDR) correction, GO Term Restriction Level 3-8 and 3 genes/4% minimum, GO Term Connection (Kappa) minimum 0.4, GO Term Grouping was on, with an initial group size of 3 and Group Merging set at 50%. Only pathway with pV<0.05 were shown. Funrich v3.0 tool (http://funrich.org/index.html) with default settings was alternatively used for functional enrichment analyses.

### Real time PCR

Total RNA was extracted using TRI Reagent^®^ (Sigma-Aldrich). RNA (3 μg) was retro-transcripted by using M-MLV (Promega, Madison, WI). qPCR was performed in triplicate by using validated qPCR primers (BLAST), Applied Biosystems™ *Power*™ SYBR™ Green Master Mix, and the QuantStudio3 Real-Time PCR System (Thermo Fisher, Whaltam, MA, USA) as previously described (Aquilano et al., 2016). mRNA levels were normalized to actin mRNA, and the relative mRNA levels were determined through the 2^−ΔΔCt^ method.

### Immunoblotting

Tissues or cells were lysed in RIPA buffer (50 mM Tris-HCl, pH 8.0, 150 mM NaCl, 12 mM deoxycholic acid, 0.5% Nonidet P-40, and protease and phosphatase inhibitors). Five μg proteins were loaded on SDS-PAGE and subjected to Western blotting. Nitrocellulose membranes were incubated with anti-HSL (4107, Cell Signalling Technology), anti-p-HSL660 (4126T, Cell Signalling Technology), anti-p-HSL565 (4137P, Cell Signalling Technology), anti-UCP1 (CS-14670S, Cell Signalling Technology) anti-p-PKA substrates (9621S, Cell Signalling Technology), anti-FXN (sc-25820, Santa Cruz Biotechnology), anti-Nrf2 (sc-722, Santa Cruz Biotechnology), anti-PPARγ (sc-7196, Santa Cruz Biotechnology), anti-Tomm20 (sc-11415, Santa Cruz Biotechnology), anti-Vdac1 (sc-8828, Santa Cruz Biotechnology), anti-Tubulin (T9026, Sigma-Aldrich) primary antibodies at 1:1000 dilution. Successively, membranes were incubated with the appropriate horseradish peroxidase-conjugated secondary antibodies. Immunoreactive bands were detected by a FluorChem FC3 System (Protein-Simple, San Jose, CA, USA) after incubation of the membranes with ECL Selected Western Blotting Detection Reagent (GE Healthcare, Pittsburgh, PA, USA). Densitometric analyses of the immunoreactive bands were performed by the FluorChem FC3 Analysis Software.

### Statistical analysis

The results are presented as means ± S.D. Statistical analyses were carried out by using the Student’s *t* test to compare the means of two groups. One-way ANOVA followed by Tukey’s test was used for comparing the means of more than two groups. Differences were considered to be significant at p<0.05.

## ACKNOLEDGEMENTS

This work was supported by Friedreich Ataxia Research Alliance (General Research Grant 2017-2018) to K.A., National Ataxia Foundation (Seed Money Grant 2016) to K.A and European Foundation for the Study of Diabetes (EFSD/Lilly_2017) to D.L.B.

## CONFLICT OF INTEREST

Authors declare no conflict of interest.

## AUTHOR CONTRIBUTIONS

K.A. conceptualized and designed the study, wrote the manuscript

D.L.B. analyzed and interpreted the data, wrote the manuscript

R.T., designed and supervised the in vivo experiments and performed research

G.G., V.C., L.D.A., R.B. performed in vivo experiments and collected results

F.T., performed in vitro experiments

F.I., M.F. performed computational analyses

S.C., M.Fr. performed immunohistochemistry experiments and optical microscopy analyses;

S.C., M.Z. performed electron microscopy experiments and analyses.

S.R., S.M., M.M., M.F., R.F. contributed in interpreting the data and in planning the research.

